# Universal Microfluidic System for Analysis and Control of Cell Dynamics

**DOI:** 10.1101/157057

**Authors:** Ce Zhang, Hsiung-Lin Tu, Gengjie Jia, Tanzila Mukhtar, Verdon Taylor, Andrey Rzhetsky, Savaş Tay

## Abstract

Dynamical control of the cellular microenvironment is highly desired for quantitative studies of stem cells and immune signaling. Here, we present an automated microfluidic system for high-throughput culture, differentiation and analysis of a wide range of cells in precisely defined dynamic microenvironments recapitulating cellular niches. This system delivers complex, time-varying biochemical signals to 1,500 independently programmable cultures containing either single cells, 2-D populations, or 3-D organoids, and dynamically stimulates adherent or non-adherent cells while tracking and retrieving them for end-point analysis. Using this system, we investigated the signaling landscape of neural stem cell differentiation under combinatorial and dynamic stimulation with growth factors. Experimental and computational analyses identified “cellular logic rules” for stem cell differentiation, and demonstrated the importance of signaling sequence and timing in brain development. This universal platform greatly enhances capabilities of microfluidic cell culture, allows dissection of previously hidden aspects of cellular dynamics, and enables accelerated biological discovery.

## INTRODUCTION

Cells operate in highly dynamic microenvironments where the type and concentration of signaling molecules are ever changing. For example, the stem cell niche presents a range of signaling molecules and growth factors regulating the maintenance of stem cell pool. During development or injury, the composition of the niche rapidly changes to allow differentiation into different lineages. The signals received by cells at various decision points on the epigenetic landscape determine their differentiation trajectories (Conrad et al., 1957; Hemberger et al., 2009). Similarly, immune cells operate in tissues and mature in specialized microenvironments in the lymph nodes or the thymus, and infection leads to modification of these environments and the chemical signals therein. It is highly desirable to recapitulate such dynamic signaling environments *in vitro* for the quantitative study as well as applications in stem cells, tissue regeneration and immunotherapy.

Signaling inputs in cellular niches include the simultaneous presence of multiple ligands (combinatorial inputs), as well as the temporal presence of ligands one after another (sequential inputs) (Figure 1A) (Judge et al., 2005). Characterizing how cells respond to such combinatorial and dynamic inputs can not only help to elucidate the regulatory mechanism underlying fundamental cellular processes, but also pave the way towards developing therapeutics through organ-on-a-chip approaches. Traditional dish-and-pipette culture techniques, however, are severely limited in creating and studying such complex dynamical microenvironments and for quantification of single live cells.

**Figure 1.**
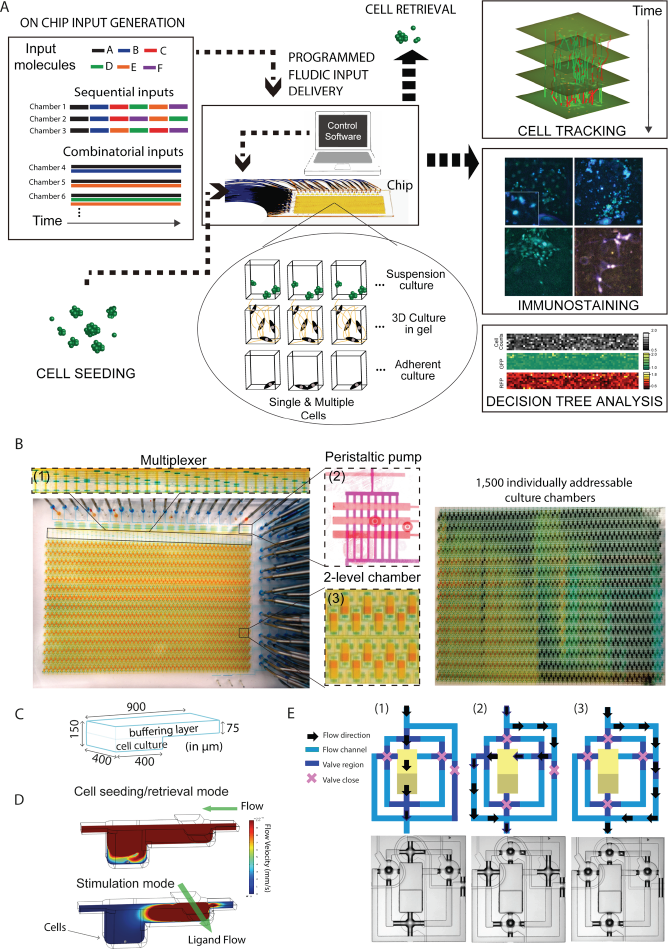
Multifunctional versatile culture system for high throughput, dynamical analysis of single-cells, cell populations and organoids. **(A)** This microfluidic system performs several automated cell culture processes such as cell seeding, stimulation, time-lapse imaging and cell tracking, and cell retrieval in a single platform, using a 2-layer culture chamber geometry and integrated PDMS membrane valves. An on-chip multiplexer measures several fluids containing signaling molecules or drugs, and mixes them at predetermined ratios, creating complex chemical inputs. A peristaltic pump with nanoliter precision delivers these inputs to 1,500 independent cell culture chambers for dynamical cell stimulation. Each of the 1,500 chambers can be programmed to receive a different input (chemical stimulus). For the combinatorial input scenario, several chemicals are mixed at a predetermined fashion and delivered to the cells, and are maintained throughout the entire experiment. In sequential inputs, signaling molecules are changed with a programmed time interval(such as 1 day, or a few minutes) to achieve dynamical cell stimulation. Both single cells and cell populations, and adherent or non-adherent cells can be cultured in suspension mode, as 2-D (monolayer), or in 3-D format using hydrogels. The system is integrated to a fluorescent microscope and can automatically track individual cells or populations in time-lapse experiments. The two-layer geometry of the culture chambers and diffusion based input/media delivery creates a stable environment for cells, allowing excellent cell viability and the ability of single-cell tracking of even non-adherent (suspension) cells during dynamical chemical stimulation. Cells can be immune-stained during or at the end of the experiments, and image processing reveals protein expression and morphology information at the single cell level, allowing quantitative computational analysis. Single cells or populations of interest can be automatically retrieved from individual chambers for off-chip analysis or expansion. **(B)** (Left) Optical image of the device containing 1,500 individually-addressable cell culture chambers loaded with orange dye in the flow layer and blue dye in the control layer. The key features of the chip are enlarged: (1) the multiplexer module, (2) a peristaltic pump equipped with 8 parallel flow channels (100 ìm in width). Fluid is driven by 3 control lines (200 um in width), (3) Shear-free cell culture chambers. (Right) Each of the 1,500 culture chambers are loaded with a different color generated through mixing of different ratios of blue, green, and red dyes, showing individual addressability. **(C)** Schematic view of the cell culture chamber composed of a buffering layer (75 μm in height) and a deeper culture unit (150 μm in height). Buffering layer is used to deliver media or signaling molecules during stimulation mode, and cell culture chamber remains flow free. **(D)** Numerical simulation of flow profile through the culture chamber during different operation modes. In the cell-loading and retrieval mode (upper panel), the flow comes from the right, and the bottom of the culture chamber experiences 1 mm/sec flow rate. Using higher flow rates allows retrieval of detached cells. During stimulation mode (lower panel), the fluid comes perpendicularly through a second channel and the flow velocity at the bottom of cell culture chamber remains zero. Cell stimulation is achieved by diffusion in this mode. **(E)** Schematic drawings (top row) and optical images (bottom row) of three distinct flow modes. (1) Fluid is directed to flow over the culture chamber directly (cell loading and retrieval mode); (2) Fluid is guided through the buffering region from the side (stimulation mode); (3) Fluid can be directed to totally bypass the chamber unit to, for example, avoid cross-contamination or perform other fluid manipulation.

In this regard, promising microfluidic cell culture technologies have been implemented to improve previously time-consuming and labor-intensive tasks into a series of streamlined and mostly automatic operations (Unger et al., 2000; Gomez-Sjoberg et al., 2007; Lecault et al., 2011; Sackmann et al., 2014), and allow realization of previously intractable experiments (Jeong et al., 2015; Junkin et al. 2016). A common pitfall, however, lies in the emergence of large numbers of microfluidic devices with each carrying distinct features to meet a narrow spectrum of specific needs (Cao et al., 2015; Occhetta et al., 2015; Sarioglu et al., 2015). Individual devices for sorting, culturing, stimulating, tracking and retrieving cells have been demonstrated, however, few systems attempt at bringing together these capabilities necessary for the study of signaling dynamics at the single-cell or population level. Temporally-varying (dynamic) stimulation and time-lapse tracking of adherent single-cells was previously demonstrated and resulted in several findings (Tay et al. 2010; Purvis & Lahav, 2013; Kellogg & Tay, 2015), however existing culture systems (microfluidic or otherwise) are not compatible with simultaneous dynamic stimulation and tracking of non-adherent cells. Furthermore, the number of experimental conditions created in previous microfluidic culture devices has been limited to less than 100 conditions (Gomez-Sjoberg et al., 2007), limiting their utility in screening a large number of conditions in exploratory signaling and drug studies. Additionally, while the success of microfluidic devices in culturing of already robust cell lines varied, maintaining viable microfluidic culture of sensitive primary mammalian cells was so far elusive. Primary cells are often rare, and are extremely sensitive to environmental conditions such as shear, flow, spatial confinement and replenishment rate of nutrients and signaling molecules. A microfluidic device that is capable of creating a large number of parallel dynamic culture conditions on wide range of cell types while exerting minimal adverse effects on the cellular microenvironment is highly desired in quantitative biology.

To address all of these limitations and to introduce much needed new capabilities in dynamical cell culture and control of cells, we developed an automated microfluidic culture system and associated integrated platform for dynamic stimulation, cell manipulation, and time-lapse microscopy. This system allows multi-mode cell culture (single cell, 2-D monolayer and in 3-D organoids) and dynamic stimulation, rapid on-chip mixing for the generation of diverse dynamic chemical inputs, as well as contain 1,500 individually addressable cell culture units for high-throughput quantitative studies on most major types of mammalian cells (Figure 1A). Each one of the 1,500 culture chambers can be programmed to receive a different set of signaling molecules, and the composition of these molecules can be changed over time in each chamber (Figure 1B). Coupled with custom software for chip control and computational data processing, the system can perform programmed delivery of thousands of formulated fluids to any designate on-chip culture unit, while monitoring and analyzing corresponding cellular responses via live cell microscopy and end-point biochemical or genetic analysis methods. This microfluidic system is a powerful platform for both fundamental and translational biomedical studies on cell signaling, and high-throughput dynamical screening of live cells with single-cell resolution.

We used this system to investigate dynamic signaling in two important areas, namely 1) Neural Stem Cell differentiation, and 2) Innate Immune pathway regulation. Our live-cell imaging experiments with single isolated fibroblast cells achieved dynamical stimulation of the NF-ĸB pathway without contribution from paracrine signaling factors, and showed that switch-like NF-ĸB activation (Tay, et al. 2010) is a cell intrinsic property. Our experiments using primary embryonic Neuronal Stem Cells (NSCs) and neuronal organoids demonstrated that NSCs proliferation, differentiation and lineage programming can be efficiently assessed at the single-cell level via tracking the expression level of self-renewal (Hes5) and differentiation (Dcx) markers in response to dynamic growth factor inputs. We systematically mapped the NSC differentiation and signaling landscape in high-throughput live-cell imaging experiments, and dissected the interactions between *in vivo* expressed NSCs regulatory ligands. We measured and computationally analyzed 1,500 combinatorial and sequential stimulation conditions, and showed how the NSCs differentiation or self-renewal can be programmed through a multi-route, time-dependent stimulation pattern of participating ligands. These experiments and subsequent computational decision tree analysis revealed a set of useful “cellular logic rules” by mapping the effects of either combinatorial or sequential signaling inputs on NSC differentiation and self-renewal.

## RESULTS

### Versatile micro-chamber for viable culture, dynamic stimulation, and specific retrieval of adherent and non-adherent cells

We designed a microfluidic cell culture chamber to cultivate a broad range of single cells and cell populations, and to create complex and dynamic signaling microenvironments (Figure 1). To accomplish this, precise control of environmental parameters such as cell density, surface properties, fluidic exchange, media and growth factor delivery, and humidity is required. Microfluidic devices using membrane valves allow automated generation of well-controlled programmable conditions mimicking *in vivo* microenvironments (Gomez-Sjoberg et al., 2007; Lecault et al., 2011; Luni et al., 2016; Tay et al., 2010). Despite the advances in integrated microfluidics, however, long-term cultivation of sensitive cell types, such as primary mammalian cells, remains challenging. Furthermore, the number of independent culture conditions (throughput) remains limited to less than 100 in a single device, still limiting the use of these systems in large-scale screening studies. Moreover, dynamic chemical stimulation of weakly adherent or non-adherent cells has not been possible when single-cell tracking and analysis is also required (Gomez-Sjoberg et al., 2007; Junkin et al., 2016). This is because exposing cells to flow during addition of chemical signals either completely removes or displaces the cells, thus obstructing single-cell tracking in time-lapse microscopy experiments. To address all of these limitations, we designed a microfluidic chamber geometry, and associated analytical device, and tested the technology’s robustness in creating 1,500 consistent micro-environmental conditions for long-term cellular studies (Figure 1B). This 3-D chamber consists of two separate sub-chambers (culture chamber and buffer chamber) dedicated to cell culture and media flow, and a set of orthogonal supply channels that allow feeding and stimulating non-adherent cells via diffusion, completely preventing cells from undesirable displacement (Figure 1C, 1D and 1E). The diffusion time for signaling molecules like TNF from the loading chamber to the cell chamber is only a few seconds, as shown by numerical simulations, allowing rapid saturation of the cell channel with signaling molecules (Video S1). This enables rapid yet gentle delivery of a wide range of dynamic chemical signaling inputs to the cells in the culture chamber with minimal distortion of the signal shape. By replenishing media and signaling molecules to cultured cells through diffusion, we satisfied the seemingly contradictory requirements of dynamic stimulation and indexed tracking of non-adherent cells in time-lapse microscopy experiments (Figure 2, Figure S1 and S2).

**Figure 2.**
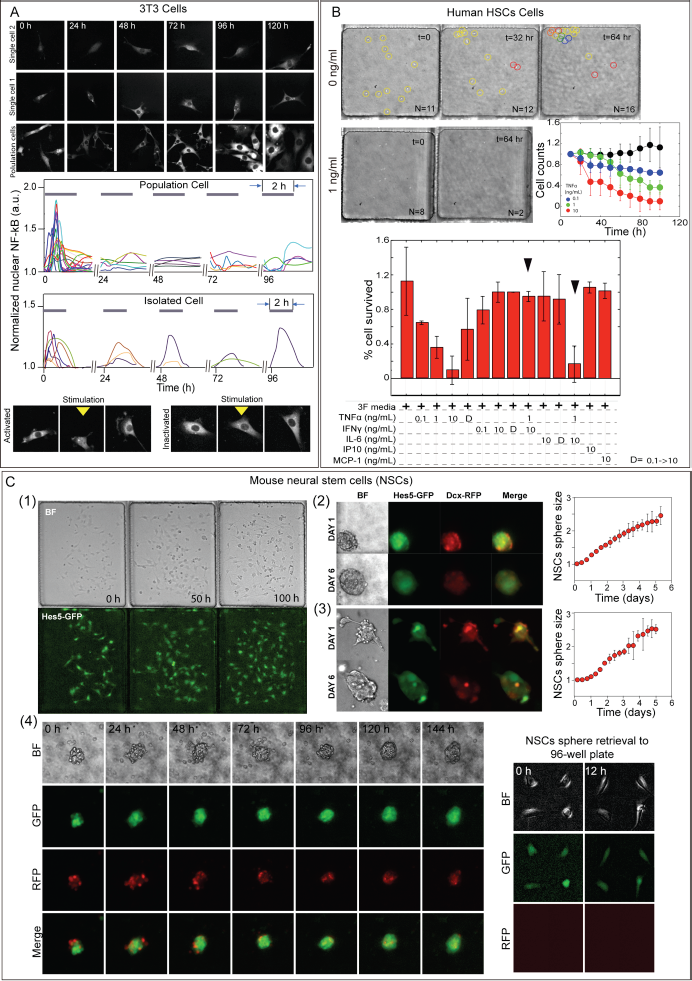
Viable and long term culture, dynamical stimulation and analysis of primary cells and cell lines, in single-cell, 2D monolayer and 3D culture modalities. **(A)** Long-term culture, tracking and biochemical stimulation of single mouse fibroblast cells in isolation or in populations. These cells express p65-DsRed for monitoring NF-KB activation. In the top panel, representative real-time fluorescent images show individual cells during 2 hour pulsed stimulation with TNF at 0.1ng/ml, in 120-hour long experiments. TNF is introduced to the designated cell culture chambers and maintained for 2 hours. After TNF stimulation, the cells are flowed with fresh media. This protocol is repeated every 24 hours. In the lower panel, representative real-time traces of p65-DsRed nuclear localization for single cells in both experiments are presented. Fluorescent images of activated and non-activated isolated single 3T3 cells at 0.1 ng/ml TNF concentration is shown. **(B)** Long-term culture and TNF stimulation of human hematopoietic stem cells (HCSs). Cells were cultured on the microfluidic device in the absence of TNF (control) and with 1ng/ml TNF, for up to 5 days. Upper right panel shows cell numbers (relative) after stimulation with TNF (black dots are control culture chambers, at [TNF]=0). N indicates cell numbers per chamber. Lower panel shows effect of TNF with the presence of other participating cytokines (IFNy, IL-6, IP10 and MCP-1). TNF induces dosage-dependent death of human HSCs, and IFNy neutralize TNF induced cell death. D represents dynamic cytokine inputs, daily variation between 0.1 and 10 ng/ml. **(C)** Long-term culture, growth and stimulation of mouse neural stem cells (NSCs) expressing key markers (Hes5 and Dcx) for self-renewal and differentiation. Three different culture modalities were used: adherent-monolayer culture, Neurosphere culture, and 3D hydrogel culture. On the left (1), we show time-lapse bright-field (top) and epi-fluorescence (GFP, bottom) images of Hes5-GFP expressing NSCs in adherent culture on chip. Hes5 expression is indicative of maintenance of the stem cell state. The time is marked at the upper-right corner of each image (in hours). In the middle-top (2), we show bright field and fluorescence images of the growth of Hes5-GFP and Dcx-RFP expressing NSCs Neurosphere in an uncoated culture chamber. Increase in Dcx expression indicates differentiation, and Neurospheres transition into a mixed population. In the middle-bottom (3), we show NSCs maintained in 3D hydrogel culture created in the chip. The corresponding growth curves for both sphere and 3D hydrogel culture are shown in the right panel. (4) Real time tracking of a NSCs sphere in a 144-hour stimulation and culture experiment. We observe that NSCs cells that express high level of Hes5-GFP signal maintain and proliferate over weeks, which is further verified by designated NSCs sphere retrieval and continue culture in 96-well plate.

The deep, well-like structure of the culture chamber allows loading and immobilizing cells by gravity, and enables growing cells into three-dimensional (3-D) colonies and organoids like Neurospheres (Figure 2C). The culture chamber can be loaded with hydrogels and other matrices to enable 3-D organization of cells, and these cells can be stimulated by providing signaling molecules through the buffer chamber, which diffuse through the gel and reach cells (Figure 2C). Furthermore, one of the supply channels can be used for on-demand retrieval of cells from the chamber and ultimately from the microfluidic device at the end of stimulation and imaging experiments, by simply increasing the flow rate in this channel (Figure 1D, 1E, 2C and Video S1). Both adherent and non-adherent cells can be automatically retrieved from the chip with high efficiency. In the case of adherent cells, treatment with trypsin is sufficient to release them from the surface (Video S2). These capabilities provide seamless and automated loading, feeding, culturing, stimulation, single-cell tracking, and retrieval of both adherent and non-adherent cell types that were maintained in isolation, as 2-D cultures or as 3-D organoids (Figure 1A). These important capabilities were never before demonstrated in a single culture system.

Our study showed that providing cells with a constant supply of fresh media at a low rate is crucial for their viability in the microfluidic environments, which we achieved here using diffusion. Due to increased surface-to-volume ratio in a microfluidic chamber, rapid water evaporation and subsequent increase in salt concentration can harm cell viability. This issue was previously addressed by introducing a large media reservoir into a third microfluidic layer (Lecault et al., 2011). Here, we can supply predetermined amounts of media to our buffer chamber, and because we completely eliminated flow from the culture chamber, many crucial culture parameters such as the humidity, CO_2_ and cell-secreted factors can be stabilized without the need for a separate media bath. Elimination of the additional PDMS membrane and media bath simplifies the chamber design, which allowed us to parallelize such independently controlled chambers and build thousands of them in a single device.

Figure 1B shows actual pictures of our microfluidic device, consisting of 3 key components: (1) a multiplexer module, (2) an array of peristaltic pumps and (3) 1,500 individually addressable flow-free 3-D culture chambers. The chamber is composed of two parts: a buffering unit (900μm × 400μm × 75μm) and a culturing unit (400μm × 400μm × 75μm, Figure 1C). During cell seeding, cells are allowed to settle to the bottom of the culture chamber while flowing slowly over the top driven by a peristaltic pump, or directed to designated culture chambers through pressure-driven flow, which is considerably faster. Four flow channels with 25μm in height and a width of 100μm allows two orthogonally controlled flow modes: top-down as the loading and stimulation mode whereas left-right as the feeding mode (Figure 1E). Numerical simulation shows that the flow in top-down direction induces a maximum flow rate of 0.6 mm/sec over cells situated at bottom of the culture unit at 1 mm/sec flow rate (Figure 1D, Cell seeding/retrieval mode). This way, culture medium can be replaced in ~5 seconds, enabling very rapid cell stimulation if needed (Video S1). Maintaining the same flow rate while directing the flow through the buffering layer (Figure 1D and 1E, Stimulation mode) reduces the flow rate at the bottom of the culture chamber to zero. Through diffusion, approximately 80% of protein-sized molecules in the culture chamber can be replaced within 20 seconds, which is sufficiently fast for many biological signaling process that operate in much slower time scales (Video S1) (Kellogg et al., 2015; Kholodenko et al., 2006; Kholodenko et al., 2010; Tay et al., 2010; Oudenaarden et al., 2013; Purvis and Lahav, 2013; Sprinzak et al., 2010). Importantly, cells experience zero shear flow even at 10 mm/sec during diffusion stimulation mode. This feature makes the culture condition more consistent and robust against any geometrical differences from chamber to chamber, and chip to chip. Moreover, cells can remain in their conditioned medium (i.e. cell secreted factors) longer than flow-based microfluidic devices, preventing detrimental effects of frequent media change on cell viability and morphology (Ghasemi-Dehkordi et al., 2015; Ludwig et al., 2006; Mehling and Tay, 2014; Ying et al., 2003). These novel features allow on-demand, fast and frequent chemical stimulation with minimal disturbance to the cellular microenvironment, and provide a stable medium composition for long-term culture of a wide range of sensitive single-cells or cell populations (Figure 2).

### Integrated culture system for high-throughput, automated stimulation and analysis of cellular dynamics

The simplicity of our multi-purpose culture chamber allows high throughput integration and control, allowing testing of thousands of culture conditions. We have integrated 1,500 such chambers in a single microfluidic device, where each chamber is independent from the others and can be exposed to a different culture condition (Figure 1B). These conditions consist of different cell types, cell densities, support matrices, as well as 1,500 precisely formulated combinations of chemical inputs such as signaling molecules, growth factors, and drugs. The chemical inputs and their flow rates in each chamber can be changed over time using on-chip membrane valves and dedicated fluidic networks, further increasing the multiplexing capability and allowing dynamical stimulation. With this device, we surpassed the throughput of previous automated microfluidic cell culture devices by 15-fold in terms of distinct culture conditions generated (Gomez-Sjoberg et al., 2007).

### On-chip formulation of complex and time-varying chemical inputs

To fully utilize the high-throughput nature of this culture system, and to dynamically change the signaling microenvironments, on-chip formulation of thousands of chemical inputs is needed (Figure 1A). Previous automated culture devices typically achieved input generation by 1-to-1 combinations of externally formulated fluids, severely limiting the number of mixtures that can be used and their concentration range. To address this limitation, we designed and integrated a microfluidic chemical formulator system consisting an array of peristaltic pumps and fluidic inlets, to formulate complex and dynamic chemical inputs on demand (Figure 1B). To efficiently deliver a formulated solution to any of the 1,500 designated culture units, we upgraded the peristaltic pump by integrating 8 parallel channels (100μm in width). The enhanced pump can achieve approximately 30 nl/sec at 10 Hz pumping frequency and thus ensure rapid and precise delivery of formulated solution. With such features, an arbitrary range of time-varying chemical inputs with distinct characteristics, such as pulsed and sinusoidal input delivery, can be generated from a few previously prepared fluid vials connected to the chip, with the temporal resolution ranging from seconds to hours. To dynamically control this microfluidic cell culture system and to automate processes like cell loading, input formulation, and fluidic delivery, we have developed specialized software. We integrated this microfluidic culture system to an automated fluorescent microscope to implement a total-analysis system capable of monitoring single cells and cell populations via live-cell microscopy in weeks-long experiments in precisely formulated signaling microenvironments (Figure 1A). The combination of on-chip dynamic input formulation, 1,500 independent culture chambers, diffusion based stimulation, and the ability to culture in all relevant modalities (single cell, 2-D or 3-D) results in the elimination of most major limitations of microfluidic cell culture, and greatly enhances its capabilities for applications when dynamic control of cells is required.

### Viable, high-throughput culture of primary single-cells and 2-D cell populations

The Achilles heel for automated microfluidic cell culture has been the poor viability of cells grown in microchambers, which is especially severe for sensitive primary mammalian cells (Mehling and Tay, 2014; Millet et al., 2007). To demonstrate the capability of our system in viable culture and stimulation of different cell types (both cell lines and primary cells), we first cultured a mouse fibroblast 3T3 cell line and primary human hematopoietic stem cells (HSCs) in adherent and suspension culture modes, respectively, and stimulated them either with constant, pulsed or sinusoidal formulations of the inflammatory cytokine TNF to induce NF-ĸB signaling (Kellogg et al., 2015; Tay et al., 2010) (Figure 2A). Dynamic (i.e. pulsed or sinusoidal) stimulation and indexed tracking of non-adherent cells like HSCs was not feasible before our study. The importance of NF-ĸB gene network resides in its role in integration of dynamic signals from a multitude of extracellular sources; this network is also central to regulating innate and adaptive immunity (Hoesel and Schmid, 2013; Wu and Zhou, 2010; Hoffmann et al., 2007; Nelson et al., 2004; Tay et al., 2010; Kellogg and Tay, 2015).

Besides being a signaling molecule involved in immunity, TNF is widely employed *in vitro* to mimic the post-transplantation signaling environment (Bogunia-Kubik et al., 2003; Brown et al., 2004; Burt et al., 2017; Mohabbat et al., 2012; Wollin et al., 2009). To mimic hematopoietic stem cell transplantation (HSCT) in our culture system, we stimulated human HSC’s with different doses of TNF as well as with combinations with other cytokines (IFNγ, IL-6, IP10 and MCP-1), and with dynamic (time-dependent) variations of selected cytokines. We observed that TNF stimulation caused death of HSCs, and this effect of TNF can be eliminated by combinatorial stimulation with IL-6 (Interleukin 6), a signaling molecule that acts as both a pro-inflammatory cytokine and an anti-inflammatory cytokine depending on context (Figure 2B). The capacity of our device as a universal culture system for various cell types is further demonstrated by the culturing of suspension cells including Jurkat T cell line and mouse HSCs (Figure S2). In both cases, cells proliferate at similar, if not higher, rate than those in bulk experiments in traditional culture dishes.

### Long-term dynamic stimulation and tracking of isolated cells reveals NF-ĸB characteristics intrinsic to single-cells

Long-term culture and tracking of isolated cells is challenging, and NF-ĸB dynamics was never studied in completely isolated single cells before (Heltberg et al., 2016; Kellogg and Tay 2015; Tay et al., 2010; Ashall et al., 2009). Dynamical analysis of single cells in populations previously showed that upon TNF stimulation, the NF-ĸB pathway activates in a switch-like fashion, where cells either fully activate or ignore the inflammatory TNF signal (Heltberg et al., 2016; Tay et al., 2010). Paracrine signaling from neighboring cells is a possible mechanism for the switch-like activation, however the inability of measuring NF-ĸB dynamics in completely isolated single cells prevented the investigation of this possible mechanism. Here, we took advantage of favorable characteristics of our culture system and studied TNF-induced NF-ĸB activation in single isolated 3T3 fibroblasts and in populations under dynamical input signals.

NF-ĸB activation is characterized by the nuclear translocation and following nuclear-cytoplasmic oscillations of the transcription factor p65, and is visualized through fluorescence imaging of 3T3 mouse cells expressing p65-DsRed fusion protein (Figure 2A and Figure S1) (Kellogg et al., 2014). Both isolated single cells and populations were successfully maintained in our culture chambers for up to a week, and were stimulated with series of TNF concentrations (0.1 to 0.5 ng/ml), either constantly, or in dynamical manner using pulsatile and sinusoidal inputs (Figure 2A and Figure S1). These fluidic inputs were formulated on the chip. We found in population experiments performed on our chip that the dose dependent fraction of activated cells versus TNF concentration is consistent with previous results (Figure S1A) (Tay et al., 2010). Stimulation of isolated single cells eliminates paracrine signaling and other population effects, allowing us to study behavior intrinsic to single cells. We found that isolated cells also showed switch-like NF-ĸB activation, therefore overruling the necessity of paracrine contribution for this important signaling feature (Figure 2A, Figure S1, and Video S3). At 0.1ng/mL TNF stimulation, 56% of isolated single cells responded, while the remaining cells ignored the TNF signal, and the fraction of responding cells increased with increasing TNF dose (Table S1). Furthermore, we found that isolated single cells also show secondary NF-ĸB oscillations, indicating that paracrine signaling is not required for dynamical oscillations of NF-ĸB (Figure S1D) (Kellogg et al., 2015).

Despite these characteristics intrinsic to isolated single cells, we have seen that the population context influences certain signaling characteristics: the fraction of responding cells has increased to 90% when the culture chambers contained more than one cell at the lowest doses (Figure S1B). This shows that while switch-like NF-ĸB activation is an intrinsic property at the single cell level, the population context (i.e. existence of neighboring cells) increases the probability of individual cells’ responding to low dose TNF signal. The response time of NF-ĸB activation (time from beginning of TNF stimulation to the NF-ĸB nucleus localization peak) also depended on population context: cells in populations responded faster on average than isolated cells in all three TNF concentrations we tested (Figure S1C). To further demonstrate the suitability of our culture system in studying NF-ĸB signaling in a highly dynamical micro-environment (Kellogg and Tay, 2015), a sinusoidal TNF signal with a maximum of 5 ng/ml, and a periodicity of 150 min was generated and delivered to designated culture chambers (Video S4). We found that cells responded in an un-entrained fashion to this input, showing increased variability in NF-ĸB nuclear localization peaks recorded compared to short-pulsed repeated stimulation (Figure S1E and S1F).

In summary, these results highlighted the utility of our system for dynamical stimulation and analysis of cells in isolation and in populations, and for answering important questions in innate immune signaling.

### Viable culture, differentiation and tracking of Neural Stem Cells and Neurospheres

The 150μm height of our culture chamber allow organization of dissociated cells into 3D structures and organoids, or direct introduction of pre-formed organoids into the chip. We can also introduce hydrogels into the culture chambers. Using diffusion feeding that prevents exposing cells to media flow, we can create stable culture conditions (i.e. self-secreted factors are not washed out) and the integrity of hydrogels can be maintained. We cultured mouse embryonic primary Neural Stem Cells (NSCs) in our system in various culture modalities. Unlike mouse HSCs, which have been studied rather extensively as a model system, NSCs are extremely sensitive to their environment and there have been only limited successes for their culture in microfluidic devices (Cordey et al., 2008). Previous studies show that variations in environmental conditions may lead to spurious NSC cell response, decreased growth rate and even cell death (Wolfe and Ahsan, 2013).

Here, we demonstrate the capability of our device in long-term culturing of primary NSCs (expressing Hes5-GFP) under three distinct modes: (1) suspension, (2) adherent monolayer and (3) in 3D hydrogel culture (Figure 2C). Expression of Hes5 protein, one of the downstream targets upon Notch signaling activation, is used as the marker for mouse derived NSCs self-renewal and maintenance of the stem cell state (Basak and Taylor, 2007; Basak et al., 2012; Haas et al., 2010; Sykova and Forostyak, 2013). For adherent culture, we tested several protocols, and found that NSCs neurospheres would gradually dissociate, spread, and migrate on the PDMS surface coated with poly-lysine and laminin (Video S5). Visible features such as cell morphology, Hes5-GFP level and cell growth are monitored for a week in these dissociated cells with time-lapse microscopy. Upon spreading, the overall Hes5-GFP level of single NSCs remains unchanged. During first 24-hour of continuous feeding (20 sec feeding with 30 min interval), the cell numbers roughly double (Figure 2C).

For suspension neurosphere and 3D hydrogel culture, intact neurospheres are directly loaded into untreated culture chamber either with or without collagen hydrogel respectively (Figure 2C). Through time-lapse imaging, we observed that the diameter of neurospheres roughly double (corresponding to approximately 8-time increase in cell numbers) in both cases after 6 days of culture (Figure 2C and Video S5). Despite sharing similar growth rates, NSCs spheres cultured in 3D hydrogel exhibited more extended axons and elongated shape, suggesting higher degree of cell attachment to the surrounding hydrogel. We further demonstrated that such cultured cells can be recovered and be used for further studies by retrieving intact neurospheres from the device and cultivating them in the 96-well plates (Figure 2C). Transferring NSCs spheres cultured on chip to an attachable surface reveals the sphere-forming ability of Hes5-positive cells, and thus verifies Hes-5 as a marker for NSCs stemness (Behnan et al., 2016). No evidence of Dcx positive cells is seen among retrieved NSCs spheres, further indicating lack of differentiation (Figure 2C). Altogether, these results show that our microfluidic device is suitable for culture and study of suspended or adherent cells, sensitive primary cell types, and of 3D organoids of embryonic NSCs.

### Dynamic stimulation of Neural Stem Cells using signaling molecules expressed during early brain development

The stem cell niche consists of a combination of signaling molecules and growth factors whose type and concentration change over time. To recapitulate signaling in the niche and the early transcriptional events during mammalian forebrain development, we used our system and cultured NSCs under complex combinations and time varying signaling inputs (Figures 3-7). In RNA sequencing measurements on embryonic mouse brain tissue, 6 signaling molecules were identified whose receptors are highly expressed during early mouse forebrain development, these are *PDGF, CXCL, PACAP, DLL, Jagged,* and *EGF* (ATLAS, 2017). To understand how the NSCs differentiation or self-renewal are regulated by these participating ligands and the temporal sequence of their expression, a series of combinatorial and sequential inputs of these 6 ligands are generated in our device. In total, around 800 conditions (including sequential, combinatorial and dose-dependent conditions) are generated and delivered to designate cell culture chambers that hold primary mouse NSCs (Table S2). Some cells were stimulated with different permutations of all available ligands during the entire experiment (combinatorial inputs), while others were stimulated with a different ligand every day (sequential inputs). These cells expressed Dcx-DsRed and Hes5-GFP proteins, for detection of differentiation into different brain cell lineages or maintenance of the stem cell state, respectively (Figure 3). Quantification of these reporters via time-lapse microscopy and single-cell tracking, along with the analysis of cell morphology, growth and immunofluorescence verified either stemness/self-renewal (high Hes5, low Dcx), or differentiated neurons (low Hes5 and high Dcx, and neuron-like morphology), Oligodendrocytes (low Hes5 and Dcx, oligodendrocyte-like morphology), or Astrocytes (low Hes5, and astrocyte-like morphology) (Figure 3, Table S3). We observed differences in growth rate, morphology, Hes5 and Dcx levels in the NSCs (Figure 4A and 4B). We compared these features obtained under each condition (sequential and combinatorial) to that of control experiments where NSCs were cultured in normal media, and discovered 17 distinct “cellular logic rules” describing the combinations and sequences of signaling molecules for achieving a particular differentiation outcome. These results are highlighted in the following sections and summarized in Table S2.

**Figure 3.**
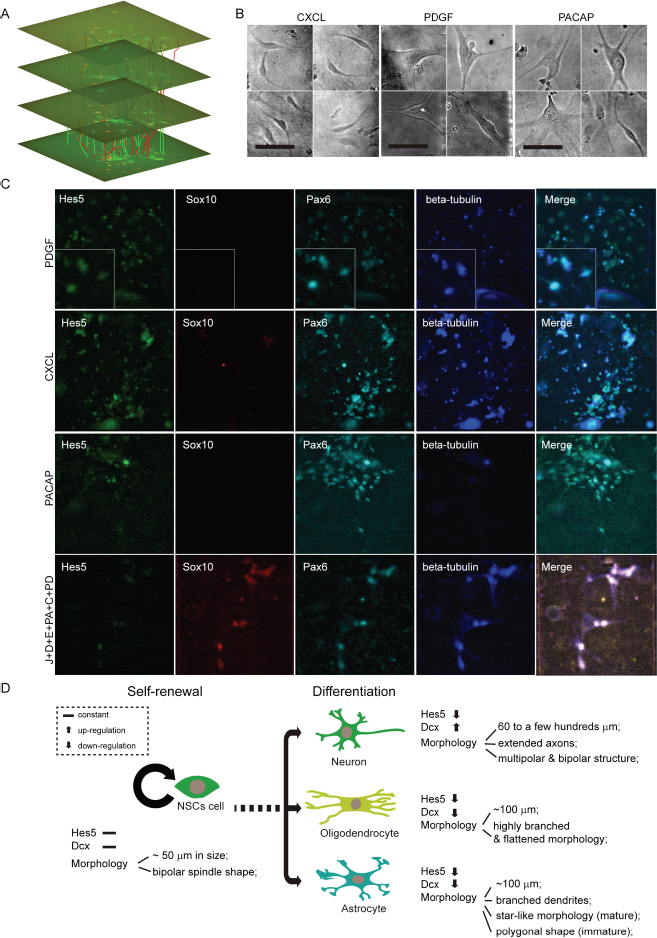
Assessment of Neural stem cell (NSC) state based on protein expression level, cell morphology and immunofluorescence in the culture system. **(A)**The temporal trajectories of NSCs maintained and stimulated in a culture chamber with a 2D+time spatiotemporal visualization (time runs along the vertical axis). The rendering also shows 4 time frames of the movie. Our device allows collecting 1,500 such videos at a time. **(B)** Bright field images of NSCs cultured on chip in media containing (from left to right) 1000 ng/ml CXCL, 50 ng/mL PDGF and 100 nM PACAP. The scale bars are 50 μm in all images. **(C)** Immunostaining images of NSCs exposed to (top row to bottom) 50 ng/mL PDGF, 1000 ng/ml CXCL and 100 nM PACAP. Markers used for determining NSCs differentiation states are Hes5, Sox10, Pax6 and beta-tubulin. Insertions in the top row are selected NSCs cells with distinct morphology. **(D)** Schematic drawing of NSCs differentiation determined by Hes5 and Dcx expression level, and cell morphology.

**Figure 4.**
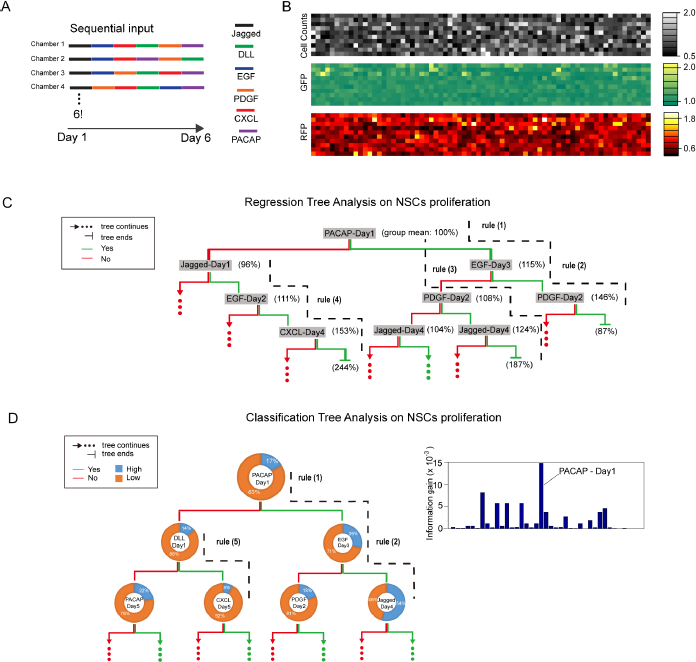
Decision tree analysis of cell proliferation under sequential stimulation reveals cellular logic rules for NSC differentiation. NSCs were stimulated with different temporal sequences of 6 ligands in the culture system, and cell numbers and Hes5 expression were quantified by time-lapse microscopy. Analysis of the cell proliferation data reveal the influence of different ligand combinations and sequences on NSC differentiation or self-renewal. **(A)** Schematic drawing of the sequential inputs composed by 6 ligands including Jagged-1, Dll-1, EGF, PDGF, CXCL-12 and PACAP-27. **(B)** Sequential inputs (720 in total) induced variations in NSCs cell counts (top), Hes5-GFP (middle) and Dcx-RFP (low) level are plotted as color maps. Squares at corresponding positions of 3 color maps represent the same experimental condition maintained in a culture chamber, and their intensities denote the relative values obtained after 6 days of stimulation. GFP and RFP intensity of individual NSCs are averaged and compared to the control experiments where NSCs were grown simply in normal media. **(C)** Regression tree analysis on NSCs’ proliferation after 6 days of stimulation. The model can be graphed as a tree-like structure, consisting of a pyramid of leaves (in grey color) that represent decision attributes - experimental conditions (a ligand tagged with its day of introduction), followed by brackets which include mean value of cell count ratio in the current group before splitting. The leaves are connected by splitting branches: the ‘Yes’ branch, i.e., the branch leading to the child set fulfilling the experimental condition, is always on the right and in green color, and the ‘No’ branch on the left and in red color. A splitting leave stops when the ratio of the current variance and variance in the starting group is smaller than 1x10^-6^, or in other words, its heterogeneity is considered to be negligible. One can read cell logic rules directly from the tree, highlighted with black dashed lines. **(D)** Classification tree analysis on NSCs’ proliferation after 6 days of stimulation. Here, each leave, plotted as a segmented ring, contains statistics on the associated dataset sorted by class labels on cell number, i.e., ‘High’ or ‘Low’, and a decision attribute at center for the splitting with its testing consequences denoted by ‘Yes’ (green) or ‘No’ (red) branches. The tree keeps growing until all instances belong to homogeneous subgroups, Information Gain approaches to zero, or the size of remaining child set is already smaller than a pre-determined number (5). Selected rules are highlighted with black dashed lines. In the right panel, Information Gain of each participating ligand with a time coordinate is plotted: *PACAP* - Day 1’s is identified to be highest and thus is chosen as the decision attribute at root.

### Decision tree analysis of sequential stimulation experiments reveals “cellular logic rules” for Neural Stem Cell differentiation

To investigate the roles of various signals and their temporal ordering in regulating NSC differentiation, we cultured and measured NSCs single cells and spheres under sequential application of six regulatory ligands (*Jagged, DLL, EGF, PACAP, CXCL,* and *PDGF)* in total 1,440 parallel culture experiments (Figure 1A, Figure 4 and 5). For each independent NSC culture, we introduced one ligand from the list each day, for a total of six days during experimental treatment. We measured: (1) The ratio of the number of cells (or the NSCs sphere size) in experiments with ligand treatment compared to the untreated control; (2) The ratio of the average fluorescence intensity of Hes5-GFP in treatment to the control experiments (Figure 4B). We subjected the resulting 1,440 experimental outputs (including both NSCs cell proliferation and Hes5/Dcx level) to regression and classification tree analyses. Seventeen distinct “cellular logic rules” leading to either differentiation or self-renewal are identified during NSCs monolayer culture, described below.

**Figure 5.**
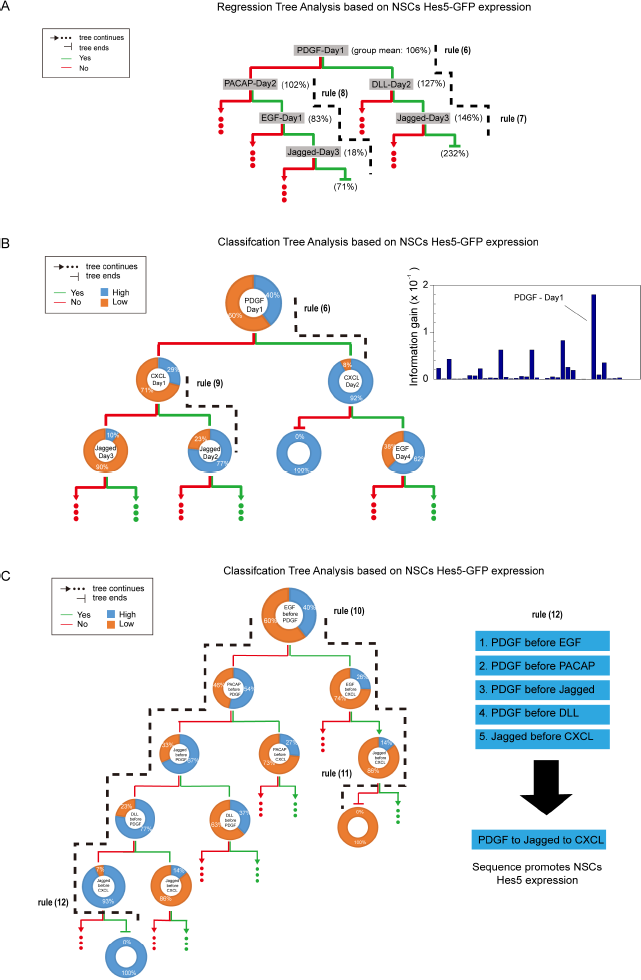
Decision tree analysis of Hes5 expression under sequential simulation reveals cellular logic rules for NSC differentiation. NSCs were stimulated with different temporal sequences of 6 ligands in the culture system, and cell numbers and Hes5 expression were quantified by time-lapse microscopy. Analysis of the Hes5 expression in live cells reveal the influence of different ligand combinations and sequences on NSC differentiation or self-renewal. **(A)** Regression tree analysis based on NSCs’ Hes5-GFP expression after 6 days of stimulation. The notations are the same as Figure 4C. Black dashed lines indicate the path leading to groups with homogenous Hes5-low or Hes5-high subgroups. **(B)** Classification tree analysis based on NSCs’ Hes5-GFP expression after 6 days of stimulation. The notations are the same as Figure 4D. Information Gains of all decision attributes are plotted in the right panel: *PDGF* - Day 1 stands out and is selected for splitting at root. **(C)** Classification tree analysis based on NSCs’ Hes5-GFP expression after 6 days of stimulation. Instead of adding simple time tags, comparative conditions (such as whether *EGF* was added before *PDGF)* are applied to categorize the whole dataset (shown at the center of segmented rings). In the right panel, by summarizing along the leftmost dashed line (rule 12), the sequence *PDGF* to *Jagged* to *CXCL* is found to promote Hes5 expression.

Using the cell count ratio as class labels, we generated decision tree models equivalent to a large number of concrete protocols for achieving particular differentiation outcome. To simplify the process, we categorize NSCs cellular states into two large groups: Cell differentiation with decreased Hes5 level and cell number; and NSC self-renewal with lifted Hes5 level and cell number. Certain input sequences stood out from numerous other possibilities. For example, cell proliferation can be preferentially triggered with any of the following sequences of signaling factors (see Figure 4C and 4D):

(1) *PACAP at day 1*
(2) *(PACAP at day 1) and (EGF at day 3)*
(3) *(PACAP at day 1) and (PDGF at day 2) and (Jagged at day 4)*
(4) *(Jagged at day 1) and (EGF at day 2) and (CXCL at day 4)*

Regression and Classification tree analysis both find rules (1) and (2). The ligand *PACAP* emerged as a dominant factor triggering cell proliferation in rules (1-3), but its presence is not absolutely necessary to increase cell proliferation, as seen in rule (4) (shown in Regression tree, Figure 4C). Rules (2) and (3) are not redundant with rule (1), as they lead to distinct rates of cell proliferation. Ligand treatment triggers increase of average cell count relative to the control (un-stimulated) sample by 146 percent and 187 percent for rules (2) and (3), respectively, compared to 115 percent and 244 percent for rules (1) and (4).

Besides, our approach identified rules that generate qualitatively distinct cellular phenotypes, such as cell differentiation, illustrated in the Classification tree as shown in Figure 4D:

(5) *DLL at day 1* ⟹ differentiation

Not only discovering quantitatively different cellular behaviors following sequential ligand administration, our approach also identified an additional collection of rules that generate qualitatively distinct cellular phenotypes, such as cell self-renewal vs. cell differentiation. Using Hes5-GFP expression ratios in a similar regression and classification tree analysis, we were able to confirm the most informative condition rule (6) in both decision trees, and discover new rules, such as rule (7-8) in Regression tree (Figure 5A) and rule (9) in Classification tree, described below (Figure 5B). In addition, using the precedence of ligand introduction as decision attributes, Classification tree found rule (10-12), as shown in Figure 5C. Here, “not” before a logical condition indicates logical negation:

(6) *PDGF at day 1* ⟹ self-renewal
(7) *(PDGF at day 1) and (DLL at day 2) and (Jagged at day 3)* ⟹ self-renewal
(8) *(EGF at day 1) and (PACAP at day 2) and (Jagged at day 3)* ⟹ differentiation
(9) *CXCL at day 1 ⟹* self-renewal
(10) *EGF before PDGF* ⟹ differentiation
(11) *(EGF before PDGF) and (EGF before CXCL) and not (Jagged before CXCL)* ⟹ differentiation
(12) *not (EGF before PDGF) and not (PACAP before PDGF) and not (Jagged before PDGF) and not (DLL before PDGF) and (Jagged before CXCL)*⟹ self-renewal

These rules show that the proliferation or self-renewal of NSCs heavily depend on the order of the signals they receive. For example, the NSCs populations almost doubled if they received *Jagged-EGF* or *PACAP-PDGF* on the first two days, and *CXCL* or *Jagged* on the fourth day (Figure 4C). Furthermore, the signals received during only the first three days of the treatment has determined Hes5 expression in NSCs (Figure 5A and 5B). The concrete cellular logic rules as well as qualitative observations emerging from the high-throughput exploration of the possible input space for NSC differentiation show the importance of individual signaling events and particularly their temporal ordering during the *in vivo* development of brain tissues. These results also constitute useful guidelines, or rules of thumb, for *in vitro* NSCs culture and differentiation experiments.

### Decision tree analysis of combinatorial stimulation reveals synergistic and inhibitory signaling interactions in NSCs differentiation

In addition to being exposed to signaling inputs in a sequential manner, cells may receive simultaneous signals from a combination of environmental stimuli. Using rapid on-chip mixing of ligands to mimic such micro-environments, we administered various constant combinations of six ligands in our device for six consecutive days, generating a total of 56 conditions (excluding single, null and all-included combinations of the six ligands) to examine possible signaling synergies among ligands (Figure 6 and Table S4). Using cell count ratio to define “high” and “low” class labels, we built a classification tree and identified another collection of rules, of which the following two are listed as an example (Figure S4A):

(13) *PACAP* ⟹ self-renewal
(14) *(not PACAP) and (PDGF and DLL)* ⟹ differentiation

**Figure 6.**
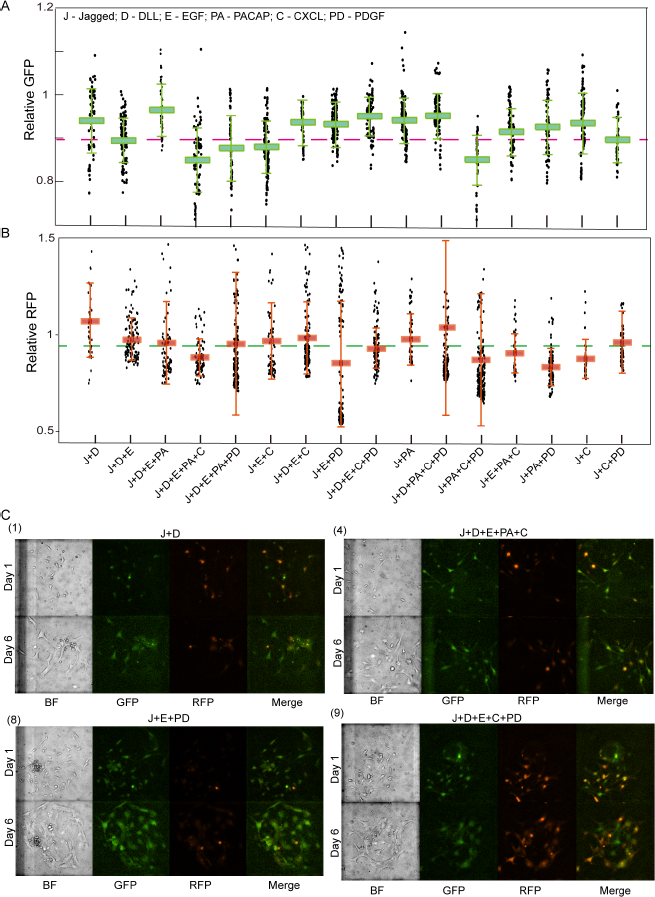
Neural Stem Cell response to combinatorial stimulation by signaling molecules that are expressed during mammalian forebrain development. **(A)** and **(B)**. Boxes represent the relative expression level of Hes5-GFP and Dcx-RFP in single NSCs, indicating stem cell status (high Hes5) or differentiation (high Dcx). Rectangular bars and solid lines represent the average and the standard deviation of all cells housed within the same chamber. Individual dots represent values obtained from each single cell, showing single-cell variability. Experimental conditions are listed in the x-axis. Each culture is stimulated with a different combination of 6 ligands that are applied continuously. Corresponding NSCs images are shown in Figure C, Supplementary figure 6 and 7. While most conditions lead to increase of Hes5 and Dcx expression, certain conditions lead to a decrease, and the data shows synergistic or antagonistic interactions between input molecules. **(C)** Selected bright-field and fluorescent images of NSCs cells on 1st day and 6th day with selected combinatorial inputs: (1) Jagged and DLL; (4) Jagged, DLL, EGF, PACAP and CXCL; (8) Jagged, EGF and PDGF; (9) Jagged, DLL, EGF CXCL and PDGF. The images are marked according to the order presented in Figure (A) and (B).

As shown in rule (13), we found *PACAP* to be the dominant factor in promoting NSCs cell proliferation, in the presence of which the percentage of high-cell-number favorite conditions increases from 38 percent to 53 percent. Rule (14) above demonstrates that without *PACAP’s* pre-existence, *PDGF* and *DLL* act synergistically to promote cell differentiation. Interestingly, the same ligand produces distinct outcomes when present at different positions of the tree. This suggests that the ligand’s function can vary according to pre-applied treatments, i.e. the signaling history of the cells. For instance, the presence of *PDGF* may lead to a pure group of either low or high cell number depending on whether *EGF* was pre-applied (highlighted by dashed lines around *EGF* in Figure S4A). Finally, it is also of great interest to investigate whether the dominance of *PACAP* can be alternatively achieved by combination of other two ligands. To that end, we subsequently found that five synergistic pairs *(Jagged + EGF, Jagged + CXCL, Jagged + PDGF, EGF + PDGF,* and *CXCL + PDGF)* tend to inhibit cell proliferation, outperforming *PACAP* alone in affecting cell proliferation (see Table S5 for their Information Gains).

Applying the same procedure to Hes5-GFP labeled dataset, we found additional rules (Supplementary Figure 4B):

(15) *PDGF* ⟹ self-renewal
(16) *PDGF and not DLL and not CXCL* ⟹ self-renewal
(17) *not PDGF and not DLL* ⟹ differentiation

Rule (15) shows the percentage of Hes5-high population increases from 39 percent to 53 percent with the presence of *PDGF,* but down to 23 percent otherwise, highlighting *PDGF’s* dominant role in promoting self-renewal of NSCs marked by high Hes5 expression (Basak and Taylor, 2007; Ohtsuka et al., 2001). Furthermore, adding *PDGF* but without *DLL* or *CXCL* (which leads to a pure Hes5-high subgroup), helps to strongly maintain NSC self-renewal (rule 16). On the other hand, adding ligands without *PDGF* or *DLL* ends up with a pure Hes5-low subgroup, resulting in cell differentiation (rule 17). Finally, out of 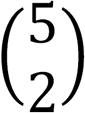possible combinations, two pairs *(Jagged + DLL* and *Jagged + PACAP)* can act synergistically to match *PDGF* alone in promoting cell self-renewal (see Table S5 for the associated Information Gains).

## Discussion

The microfluidic system described here represents the next generation for cell culture and live-cell analysis systems, suitable for a wide range of applications in stem cells, organoids, signaling dynamics, and high-throughput screening. We introduce several technical advances in this study, leading to significant improvement of the capabilities of microfluidic or other culture systems in several aspects: First, our integrated device uses diffusion instead of flow for media and signal delivery to both adherent and non-adherent cells, while maintaining a stable cellular microenvironment. The capacity of delivering both static and dynamic chemical inputs (such as pulsed or sinusoidal), together with 1,500 individually programmable cell culture units, provides powerful solutions for live-cell screening studies, particularly for the studies of the complex environmental effects on cellular responses, most of which used to be technically very challenging and time-consuming before this study.

A key advantage of our two-layer culture chamber geometry is the spatial separation between the cells and the buffering area used for media delivery, protecting cells from detrimental effects of flow (Figure 1C and 1D). In contrast, previous automated microfluidic cell culture chips adopt a configuration where either the fluid is directly flowed over the top of cells and tissues or an additional membrane is needed to isolate biological samples from flow (Gomez-Sjoberg, et al., 2007; Lecault, et al., 2011; Woo, et al., 2009). Membrane barriers isolate shear flow from the biological samples. Their application, however, is limited by the molecular diffusion rate through the membrane barriers, slowing down signaling input delivery, and by the technological challenges in fabricating such membranes in high-throughput devices. Although traditional micro-wells can culture primary cells and deliver fluids using manual or robotic pipetting, the shear force exerted on cells is non-negligible, and prevents tracking of an important class of cells that do not adhere. The 2-level device reported here provides a simple yet capable solution with an integrated buffering layer that serves as a cushion during media exchange. We found that there is no fluid flow at the bottom of the culture unit, for up to 10 mm/sec flow rate in the buffer chamber (Figure 1D and Video S1). This robustness against shear is critical for studies on sensitive primary cells and other delicate biological samples, and especially for time-lapse study and tracking of non-adherent cells. Dynamic stimulation and real-time tracking of non-adherent or weakly adherent cells was not possible in previous microfluidic culture systems. While maintaining cells in the shear-free environment, our device is able to perform rapid and complete fluid exchange, either by direct flow or by diffusion (Video S1). With 4-valve control, fluids can be directed either through the buffering layer or directly over the culture chamber (Figure 1D and 1E). Additionally, the relatively large (150μm) height of the culture chamber further extends the chip’s functionality to accommodate biological samples in 3D micro-environment such as embryonic NSCs neurospheres or other organoids (Figure 2C).

Our “Universal Cell Culture Chip” viably cultures, dynamically stimulates and tracks a wide range of cell lines, both adherent and non-adherent cells, and primary cells and cell lines. We demonstrated culture of fibroblast 3T3, Raw 264.7 and Jurkat T cells, and primary cells such as mouse HSCs, human HSCs and mouse embryonic NSCs. We found that in our device fibroblast 3T3 and Jurkat T cells proliferated at a rate comparable to that in a traditional culture in a dish (Figure S2A). Automated cell retrieval from the chip using 3T3 and Jurkat cells achieved >95% retrieval efficiency in our experiments (Video S2).

Population NF-ĸB response to TNF stimulation was seen to reproduce previously published findings in this important signaling pathway (Kellogg and Tay, 2015; Tay et al., 2010). Due to favorable conditions in our device compared to previous microfluidic culture systems, we were able to culture isolated single fibroblast cells for long durations and stimulate them with various doses and temporal impulse shapes of TNF. Such analysis of NF-ĸB pathway in completely isolated single live cells which was not demonstrated before, and it eliminates possible population effects (i.e. paracrine signaling), leading to understanding of single-cell behavior. These experiments revealed that the digital activation of NF-ĸB in single cells is a cell-intrinsic property, and does not require a contribution from paracrine signaling. On the other hand, the population context increases the probability and speed of individual cells responding to TNF input.

Our system can efficiently culture and monitor highly sensitive and rare cell types like human hematopoietic stem cells (HSCs). We observed proliferation of purified human CD34+ cells in 6-day long experiments, which are 98% pure HSCs, and have been typically used to model HSC transplant in preclinical mouse models (Figure 2B and Figure S2B). In a typical experiment, human CD34+ cells are much lower in initial cell number (a few thousands per harvest), compared to freshly isolated CD45+EPCR+CD48−CD150+ (E-SLAM) adult mouse bone marrow HSC cells, which were also studied in our system. Loading approximately 10 cells per chamber, our microfluidic device allows the study human HSCs under up to 200 conditions upon 1 sample harvest, which generally provide only a few thousand cells. The rapid response of mouse HSCs to M-CSF input suggests that media replacement through the buffering layer is effective (Figure S2B). We observed that HSCs stimulated with the pro-inflammatory cytokine TNF died in a dose-dependent manner, where higher doses of TNF led to increased HSC death (Figure 2B).

The 3-D chamber geometry of our device allows culture and stimulation in different modes, as single cells, surface bound monolayers, or 3D cultures (in gel or as suspension). With 150μm chamber height, primary neurospheres with diameters ranging from 10 to 100μm were maintained in our device and were allowed to grow. The diameter of the NSCs spheres doubles after 5-day culture on chip, indicating approximately 8 times increase in cell number (Figure 2C). In 3D hydrogel culture, despite that NSCs may adopt different morphologies, a similar growth rate is observed (Figure 2C). In surface-adherent culture, NSCs cell number doubles within the first 24 hours.

Our device incorporates 1,500 independent culture chambers, with dedicated and independent fluidic inlets and outlets, and therefore exceeds the throughput of independently addressable microfluidic cell culture systems by nearly 15-fold (Gomez-Sjoberg et al., 2007). This record-breaking throughput enables screening of thousands of dynamical input conditions in a single experiment. Single cells or cell populations can be viably cultured, dynamically stimulated with complex signaling inputs with extremely high temporal precision, and be tracked by live-cell microscopy. For off-chip analysis or expansion, cells of interest can be retrieved from the system without disturbing other cultures. On-chip integration of microfluidic peristaltic pumps, input mixing and formulation structures, and automation of all of these tasks allows a truly capable system for high-throughput imaging and analysis in quantitative biology studies.

To demonstrate the utility of our system in high-throughput live-cell imaging and quantitative analysis of primary cells, we investigated the self-renewal and differentiation dynamics of neuronal stem cells. High-throughput integration of independently programmable chambers together with on-chip formulation of chemical inputs enabled previously intractable experiments that mimic complex cellular microenvironments (including dynamic ligand inputs). As a demonstration, 6 selected ligands that bind to highly expressed receptors during early embryonic forebrain development *(PDGF-AA, PACAP, CXCL, EGF, Jagged, DLL)* were introduced into designated culture chambers either as different combinations or time sequences (Table S2 and S4). High-throughput quantitative data sets on NSCs proliferation, Hes5 and Dcx expression level, and cell morphology were obtained through live cell imaging and automated single cell tracking and analysis (Video S5). NSCs Hes5 level and proliferation rate vary in non-trivial fashion in response to static and dynamic inputs (Figure 4B and Figure 6), and their quantitative results are summarized in Table S2 and S4.

Through decision tree analysis, we discovered a set of “cellular logic rules” by mapping the effects of either combinatorial or sequential signaling inputs on NSC differentiation and self-renewal. Among these signaling molecules, *PDGF* was identified as the dominant factor causing high Hes5 expression (indicating retention of the stem cell state) while *PACAP* strongly promoted NSCs proliferation (Figure 4 and Figure S4). These ligands led to maintenance and growth of the stem cell pool. We confirmed this finding in additional experiments, in which the sole presence of ligand PDGF increases Hes5 level by around 25% from its initial value (Figure 7). The few NSCs cultures showing low growth rate and extended bipolar morphology (~100μm in size) are shown to be initially immature neurons through immunostaining (+beta-tubulin, +Hes5) (Figure 3B and 3C). Meanwhile, most NSCs under *PDGF* stimulation retained a stem cell like morphology (bipolar, ~ 50 μm), suggesting the key role of *PDGF* in NSCs self-renewal. In contrast to stimulation with PDGF only, all NSCs exposed to combination of all 6 ligands showed increased beta-tubulin and decreased Hes5 level. With morphologies carrying a bipolar structure with extended axons, the data indicates these NSCs has either differentiated or on its way to become mature neurons (Figure 3D)(Otify et al., 2014; Xiong et al., 2014).

**Figure 7.**
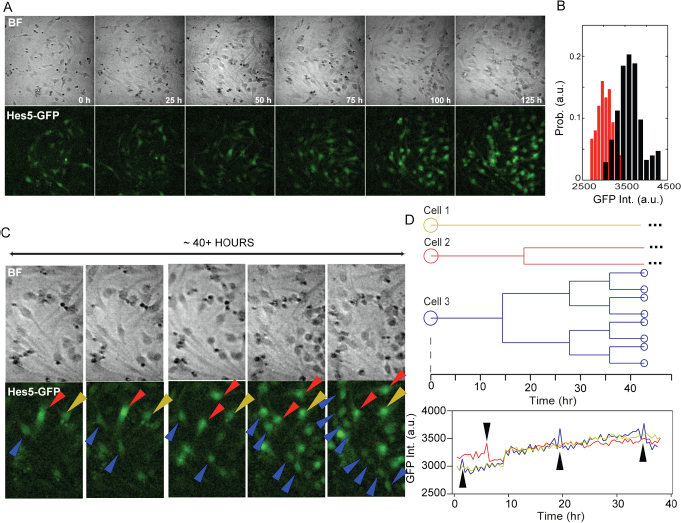
PDGF is the dominant signaling factor in NSCs self-renewal and proliferation. **(A)**Time-lapse bright field (top) and epifluorescence (bottom) images of NSCs cultured in the media containing 50 ng/ml PDGF. **(B)** Histogram of the Hes5-GFP expression of NSCs before (red) and after (black) one week incubation with the media containing 50 ng/ml PDGF. **(C)** Enlarged bright field (top) and their corresponding epifluorescence (bottom) images of Hes5-GFP NSCs cultured on chip in the media containing 50 ng/mL PDGF. 3 selected cells were indicated by differently colored arrows and tracked over 40 hours during on chip culture. Distinct proliferation pattern was observed. **(D)** Lineage tracing (top) and Hes5-GFP level (bottom) as a function of 40 hours for the 3 selected cells in Figure C.

After experimentally measuring all 720 permutations of the 6 ligands in a temporal manner, input sequences leading to high Hes5 expression and fast growth rate were identified through regression or classification tree analysis (Figure 5). The dominant role of *PACAP* or *PDGF* in promoting either NSC self-renewal or cell proliferation is further confirmed in this experimental setting: through quantitative analysis we found that the presence of *PACAP* or *PDGF* simply at 1^st^ day of a total 6-day experiment is enough to drive high cell proliferation or Hes5 level respectively (Rules 1 – 3, 6 and 7). Another notable input sequence leading to higher cell proliferation includes *Jagged* at day 1, *EGF* at day 2 and *CXCL* at day 4 (rule 4). Sequences such as *DLL* at day 1 (rule 5) as well as *EGF* at day 1, *PACAP* at day 2, and *Jagged* at day 3 (rule 8), on the other hand, was found to preferentially cause NSC differentiation.

These experimental and computational analyses attempted at mapping the signaling landscape in neuronal stem cell differentiation. By combinatorial and temporal stimulation with signaling molecules, we identified certain cellular decision points and differentiation trajectories. Interestingly, we found that many differentiation outcomes can be achieved by multiple signaling routes, showing the complex landscape and the redundancy in NSC differentiation pathways. High-throughput experiments and computational analysis enabled by our microfluidic device revealed several non-trivial signaling rules, highlighting the importance and role of the dynamically changing microenvironments in regulating stem cell behavior. The order of these *in vitro* signaling inputs made a significant difference in regulating NSC outcomes, shedding light on the in vivo differentiation processes.

In summary, the high-throughput microfluidic device shown here creates a long-term stable, shear-free micro-environment that could be dynamically modulated when needed, and enables the study of both suspension and adherent cell types under complex but pre-programmable dynamic inputs. We demonstrate its versatility and capability by successfully cultivating various cell lines as well as extremely sensitive primary cells, and by identification of signaling rules for stem cell differentiation. With its capability of efficient on-chip mixing and chemical input formulation, the device is perfectly suited for combinatorial or systematic dissection of the signaling landscape of stem or immune cells, and is immediately applicable to personalized therapy study by, for example, growing tumor organoids or immune cells extracted directly from patients and by performing drug screening. We envision that such high-throughput study will greatly shorten the time span and reproducibility in traditional screening process. Applying the same methodology, one may, for the first time, reconstruct the complex cellular environment with dynamic inputs and ultimately mimic various developmental processes of tissues and organs in vitro. With all these exciting future directions in mind, our studies on pluripotency and differentiation of stem cells (HSCs and NSCs) under dynamic conditions serve as a starting point for the next-generation organ-on-chip studies.

## EXPERIMENTAL AND ANALYTICAL PROCEDURES

### Design and Fabrication of Microfluidic Chips

We designed and fabricated the microfluidic device according to the standard protocol, which is reported elsewhere (Unger, et al., 2000). Briefly, we designed our two-layer device using AutoCAD (Autodesk Inc., San Rafael, CA, USA), and then printed the sketch on transparencies at 40 kdpi resolution (Fine Line Imaging, Minneapolis, USA). Molds for PDMS casting were produced using standard soft-lithography. The channel network of control layer as well as the flow channels for flow layer and culture chambers was produced with either SU-8 3025 or SU-8 3075 (Microchem, Westborough, MA, USA) on silicon wafers. For the flow layer, we additionally used AZ-50X (AZ Electronic Materials, Luxembourg) at valve positions. Photoresists were spun to a height of 25μm for channels, and 150μm for culture chambers. To fabricate the chip, 72g of PDMS (10 : 1 of monomer : catalyst ratio) was mixed, de-bubbled and poured over the Trimethylchlorosilane treated patterned silicon wafer. The PDMS was then cured for 60 min at 80°C. Following plasma and alignment between flow and control layer, inlet holes were then punched after two-hour thermal bonding. The chip was bonded to a PDMS coated coverslip and cured for at least 12 hours at 80 °C before use.

### Chip Setup, Operation and Control

The glass slide carrying the microfluidic chip was cleaned and taped on a slide holder. Control channels were connected to miniature pneumatic solenoid valves (Festo, Switzerland) that were controlled with a custom MATLAB (MathWorks, US) through graphical user interface (Gomez, et al). Optimal closing pressures of push-up PDMS membrane valves were determined individually for each chip, typically ranging from 25 to 30 psi. The cell culture chambers were treated with either fibronectin (0.25 mg/mL; Millipore, Austria) for 3T3 cell culture or poly-lysine (0.01%, Sigma-Aldrich) followed by laminin (1 to 2 mg/ml, Sigma-Aldrich) for adherent NSCs culture. The remaining coating solution was flushed off from the chip using either PBS or cell culture media. Cell culture media is pre-warmed on chip for at least one hour before cell loading.

### Cell Culture and Loading

For standard cell lines, we used Jurkat cells, RAW 264.7 macrophages p65-/- with p65-GFP and H2B-dsRed, as well as NIH 3T3 p65-/- cells with p65-dsRed and H2B-GFP for tracking and analysis of NF-ĸB activation. These cells are cultured according to the established protocols (Lee, et al., 2014). To seed cells into the chip, adherent cells are harvested at 80% confluence with trypsin, re-suspended and loaded into chips through semi-automated loading program at cell density from 10^4^ to 10^6^ per milliliter depending on the desired cell density.

Murine hematopoietic stem cells (HSC) are isolated by FACSAria III flow cytometer (BD Biosciences) as Lin-/ c-Kit+/ Sca-1+/ CD48-/ CD150+/ CD34- (lineage, Lin: CD3e/ CD11b/ CD19/ CD41/ B220/ Gr-1/ Ter-119), which are approximately 50% pure HSCs. Macrophage colony-stimulating factor (M-CSF), a myeloid cytokine released during infection and inflammation, is employed to induce HSCs differentiation. Human CD34+ cells were isolated from mononuclear cells using EasySepTM human CD34 positive selection kit (Stemcell Technologies, Vancouver, BC, Canada). CD34+CD38-CD45RA-CD90+CD49f+ HSCs were sorted using a FACSAria III flow cytometer (BD Biosciences). Pro- and anti-inflammatory cytokines including TNF, IFNγ, IL-6, IP10 and MCP-1 are introduced into the cellular environment as single ligand and in combination. Embryonic NSCs with Hes-GFP and Dcx-RFP reporters were isolated at embryonic day 12.5 from a transgenic mouse carrying Hes-GFP and Dcx-RFP using established protocol (Basak, et al., 2007). The resulting primary cells were verified to carry both Hes5-GFP and Dcx-RFP after isolation and allowed to grow for few passages before use in the experiments (Giachino, et al., 2009). NSCs were cultured as neurospheres in culture media (DMEM/F12 + Glutamax (Gibco No:31331-028); 10 U/ml penicillin; 10 ug/ml Streptomycin; B27 supplement (1:50); FGF growth factor 0.02 ug/ml). As NSCs are sensitive to environmental variations, cell handling protocol before loading into the chip is examined systematically (including dissociation, FACS sorting, etc.). To obtain the optimal results, NSCs spheres are collected and loaded into the chip after 24-hour culture in plate. There are, approximately, 7 to 10 cells in each sphere. To avoid potential artifacts due to prolonged in-vitro culture, only NSCs within 10 passages were used in the study.

The environmental conditions are maintained using temperature control and incubator system (Life Cell Imaging Service GmbH, Basel, Switzerland), which consists of a box surrounding the microscope, to strictly 37 °C and >98% humidity and 5% CO_2_ during the experiment, and the PDMS chip is covered with a stage-top-incubator connected to a humidifier and a gas exchanger.

### Live-cell Fluorescence Microscopy and Data analysis

For image acquisition, a Nikon Ti-ECLIPSE microscope with an automated translation stage and a digital CMOS camera (ORCA-Flash 4.0, Hamamatsu, Japan) was used. The stage and image acquisition was controlled via the NIS Elements software. Bright field and fluorescence images were acquired and analyzed using a customized MATLAB program (Mathworks, Austin, USA). The algorithm extracts single cell traces including position, nuclear and cytoplasm fluorescence level. For example, the 3T3 cell nuclear area in each image is identified via the fluorescent nuclear marker GFP, and then the mean value of the nuclear intensity of the p65-DsRed marker is measured, and plotted as a function of time.

### Decision Tree Analysis

In this work, we use decision tree analysis to generate descriptive and predictive models which map measurements about cell phenotypic features (i.e., cell count and Hes5-GFP expression) to experimental conditions (i.e., sequential and combinatorial application of six regulatory ligands). Based on the model, we can identify casual factors that determine the fate of NSCs: self-renewal and differentiation (inferred from measurements). Both regression and classification trees, as two subclasses of decision tree, were implemented. Both methods partition the data space recursively according to evolving decisions on current best split. But they handle different types of data and thus adopt different metrics for defining ‘best’ split.

Regression tree is designed for measurement of continuous or discrete ordered variables and uses Variance *Var*(*X*) as a measure of heterogeneity for the dataset X with *n* samples, which can be formulated as: 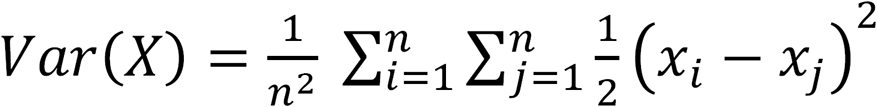. From the parental to the children sets, the goal is to reduce the variance as much as possible. We quantify the change in variance from a prior state to a state given a decision attribute α (i.e., the choice of experimental conditions) using Variance Reduction *VR*(*X, α*) as shown in the equation:

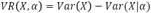

The decision attribute coming with the largest VR value is deemed to be a dominant factor. One can select the ‘best’ split accordingly, in which the reduction of variance is maximal and the most significant change in NSCs status is observed, by comparing the resulting VR values for all possible decision attributes. In this study, the decisions are True/False statements on the presence of 6 ligands - *Jagged, DLL, EGF, PACAP, CXCL, PDGF* (combinatorial inputs) and the sequence of ligand additions with time coordinates assigned (sequential inputs).

On the other hand, classification tree is typically for measurement of categorical variables, and data obtained through automated cell tracking on cell number and Hes5 levels are grouped based on two pairs of concepts: Entropy and Information Gain (IG). Different from Regression tree, entropy *H*(*X*) is used to measure the heterogeneity of the dataset and can be explicitly written as: *H*(X) = *−P*(*x_H_*) log_2_ *P*(*x_H_*) − *P*(*x_L_*) log_2_ *P*(*x_L_*), where *P*(*x_H_*) or *P*(*x_L_*) is the probability of being labelled with binary value {*high, low*}, based on pre-determined threshold (i.e., 1.4 in NSCs cell number and 1.1 in Hes5 level, both of which are ratios over baseline values in control experiments). The value of *H*(X) ranges from 0 to 1; the more heterogeneous the set is, the higher its entropy. Similarly, to reduce entropy value from the parental to the children sets as much as possible, *IG*(*X, α*), as defined in the following equation, is to be maximized given a decision attribute α:

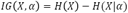

Here, ID3 (Iterative Dichotomiser 3) algorithm is adopted for decision tree analysis of NSCs data generated from combinatorial and sequential inputs (Quinlan et al., 1986).

